# The conserved herpesviral kinase ORF36 activates B2 retrotransposons during murine gammaherpesvirus infection

**DOI:** 10.1101/2020.01.06.895789

**Authors:** Aaron M. Schaller, Jessica Tucker, Ian Willis, Britt A. Glaunsinger

## Abstract

Short interspersed nuclear elements (SINEs) are RNA polymerase III (RNAPIII) transcribed, retrotransposable noncoding RNA (ncRNA) elements ubiquitously spread throughout mammalian genomes. While normally silenced in healthy somatic tissue, SINEs can be induced during infection with DNA viruses, including the model murine gammaherpesvirus MHV68. Here, we explored the mechanisms underlying MHV68 activation of SINE ncRNAs. We demonstrate that lytic MHV68 infection of B cells, macrophages and fibroblasts leads to robust activation of the B2 family of SINEs in a cell autonomous manner. B2 ncRNA induction requires neither host innate immune signaling factors nor involvement of the RNAPIII master regulator Maf1. However, we identify MHV68 ORF36, the conserved herpesviral kinase, as playing a key role in B2 induction during lytic infection. SINE activation is linked to ORF36 kinase activity and can also be induced by HDAC1/2 inhibition, which is one of the known ORF36 functions. Collectively, our data suggest that ORF36-mediated changes in chromatin modification contribute to B2 activation during MHV68 infection, and that this activity is conserved in other herpesviral protein kinase homologs.

**AUTHOR SUMMARY:** Viral infection dramatically changes the levels of many types of RNA in a cell. In particular, certain oncogenic viruses activate expression of repetitive genes called retrotransposons, which are normally silenced due to their ability to copy and spread throughout the genome. Here, we established that infection with the gammaherpesvirus MHV68 leads to a dramatic induction of a class of noncoding retrotransposons called B2 SINEs in multiple cell types. We then explored how MHV68 activates B2 SINEs, revealing a role for the conserved herpesviral protein kinase ORF36. Both ORF36 kinase-dependent and kinase-independent functions contribute to B2 induction, perhaps through ORF36 targeting of proteins involved in controlling the accessibility of chromatin surrounding SINE loci. Understanding features underlying induction of these elements following MHV68 infection should provide insight into core elements of SINE regulation, as well as dis-regulation of SINE elements associated with disease.

## INTRODUCTION

A large fraction (40-45%) of mammalian genomes is composed of sequences derived from retrotransposable elements, which are capable of copying themselves (autonomous) or being copied (non-autonomous) and inserted semi-randomly back into the genome. Retrotransposons are ubiquitously spread throughout the genome and are important components of genome architecture and chromatin remodeling [1–4]. Among these, the Short Interspersed Nuclear Element (SINE) subfamily of retrotransposons make up ∼12% of the genome and are transcribed by RNA Polymerase III (RNAPIII) to produce short ∼300bp noncoding RNAs (ncRNAs). They are evolutionarily derived from other common RNAPIII-transcribed genes, such as 7SL in the case of the human Alu SINE, and tRNA in the case of the mouse B2 SINE. SINE ncRNAs are non-autonomous and co-opt the machinery encoded by the Long Interspersed Nuclear Elements (LINEs) for reverse transcription and re-integration. SINEs may act as functional enhancers and mobile RNA polymerase II promoters, and are also present as ‘embedded elements’ in many mRNA transcripts, where they can influence mRNA processing, localization, and decay [1, 5–7].

B2 SINE ncRNA transcription is RNAPIII-dependent, requiring the transcription factor complexes TFIIIC and TFIIIB. TFIIIC binds to the internal A and B-boxes present within type-II RNAPIII promoters, such as those contained within B2 SINE and tRNA species. This is followed by recruitment of TFIIIB, comprised of BDP1, BRF1, and TBP, which help position RNAPIII at the transcription start site. Absence of BRF1 abrogates transcription from type-1 and type-II RNAPIII promoters but does not affect transcription from type-III RNAPIII promoters, which use a Brf1 paralog, Brf2 [8]. RNAPIII activity can be broadly controlled by its master repressor Maf1, a phosphoprotein that binds BRF1 and RNAPIII, thereby preventing TFIIIB assembly onto DNA and blocking the association of the polymerase with TFIIIB that is already assembled at transcription start sites, respectively [9]. Phosphorylation of Maf1, for example by mTORC1 [10], prevents Maf1-mediated repression of RNAPIII, thereby potentiating an increase in transcription.

SINE expression is normally repressed due to the maintenance of repressive tri-methylation of lysine 9 on histone H3 (H3K9me3) [11] and CpG methylation of DNA [12]. However, SINEs become de-repressed under conditions of cellular stress, such as chemical treatment and heat shock [13–15]. SINEs from both humans and mice are also induced during infection with a variety of DNA viruses, including herpes simplex virus (HSV-1), adenovirus, minute virus of mice, simian virus 40 (SV40) and murine gammaherpesvirus 68 (MHV68) [16–22]. Several recent reports indicate that virus-induced SINEs and other RNAPIII-transcribed ncRNAs interface with innate immune pathways, and thus may serve as signaling molecules during infection [23, 24]. In particular, B2 ncRNAs induced upon MHV68 infection potentiate NF-κB signaling, in part through a pathway involving the mitochondrial antiviral signaling protein MAVS, and also boost viral gene expression [21, 25]. Aberrant accumulation of Alu RNAs contributes to age-related macular degeneration by inducing cytotoxic NLRP3 inflammasome activation [26–30], and can also induce epithelial-to-mesenchymal transition, a hallmark of progression of several cancers [31]. Additionally, SINEs induced during heat shock can bind and inhibit RNA polymerase II transcription, indicating that these ncRNAs may have a variety of functions during stress [15, 32].

MHV68 is a model gammaherpesvirus related to Kaposi’s sarcoma-associated herpesvirus (KSHV) and Epstein-Barr virus (EBV), and has been widely used to dissect gammaherpesvirus biology and pathogenesis. A recent genome-wide mapping study revealed that MHV68 infection of murine fibroblasts leads to activation of ∼30,000 B2 SINE loci, although the mechanism of B2 induction is unknown [22]. Here, we show that in addition to fibroblasts, B2 SINE induction occurs during MHV68 lytic infection of primary bone marrow-derived macrophages and during lytic reactivation of B cells, both physiologically relevant cell types for the virus. Induction is cell autonomous, occurs independently of innate immune signaling components and does not involve RNAPIII regulation by the master repressor Maf1. Instead, a screen of MHV68 open reading frames (ORFs) revealed a role for the conserved herpesvirus protein kinase ORF36 in B2 SINE induction. Expression of WT ORF36 but not a kinase dead mutant was sufficient to activate B2 SINEs, and an MHV68 mutant lacking ORF36 displayed reduced SINE induction potential. ORF36 inhibits histone deacetylases 1 and 2 [33, 34] and we show that chromatin de-repression contributes to B2 activation. Collectively, our results reveal a new function for the herpesviral protein kinase and provide insight into the mechanism of SINE activation during viral infection.

## RESULTS

### MHV68 infection induces B2 SINEs in physiologically relevant antigen presenting cell types

Our previous work established that MHV68 infection of murine fibroblasts results in robust activation of B2 SINEs [21]. While fibroblasts are commonly used to study MHV68 infection *in vitro*, two of the most physiologically relevant cell types for the *in vivo* MHV68 lifecycle and establishment of lifelong latency are B cells and macrophages [35]. We therefore sought to determine whether B2 SINE induction is also a feature of MHV68 infection in these key cell types.

Although B cells are the main viral reservoir *in vivo*, they are highly resistant to de novo MHV68 infection in cell culture [36]. The only latently infected B cell line isolated from an MHV68-infected mouse tumor, S11, reactivates to very low frequency, making study of lytic cell populations impractical [37]. However, a B cell line has been generated (A20-HE-RIT) that is latently infected with MHV68 and contains a doxycycline (dox)-inducible version of the viral lytic transcriptional activator gene RTA. Treatment of these cells with Dox and phorbol ester (PMA) enables the switch from latency to lytic replication in approximately 80% of the cells [38, 39]. Induction of the lytic cycle by dox and PMA treatment of the A20-HE-RIT cells caused a marked increase in B2 SINE levels as measured by primer extension, with levels peaking at 24-32 hours post stimulation (Fig 1A). Importantly, B2 RNA induction was not seen in the uninfected A20 parental cells subjected to the same dox and PMA treatment. Furthermore, the induction observed in infected cells is specific to B2 SINEs, as levels of another RNAPIII transcript, 7SK, remained unchanged. Similar to our observations during MHV68 lytic replication in fibroblasts [21], PAA treatment to block viral DNA replication did not prevent B2 SINE induction during reactivation in A20-HE-RIT cells, although the levels were modestly reduced (Fig 1B). Thus, upon lytic reactivation of latently infected B cells, B2 SINEs are induced early in the viral lytic cycle, and continue to accumulate as infection progresses.

**Figure 1:**
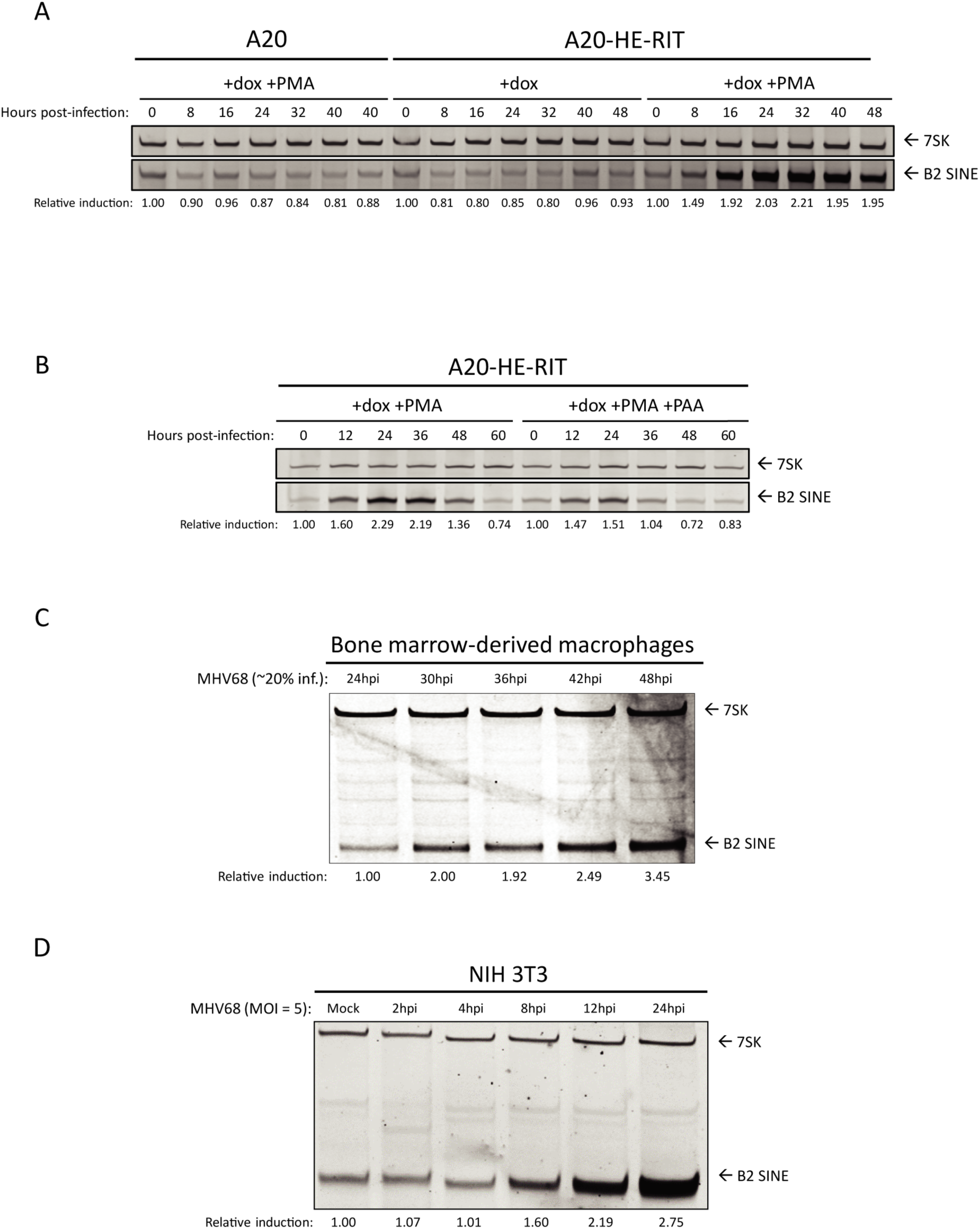
B2 SINE transcription is upregulated in B cells, primary macrophages, and NIH 3T3 cells upon MHV68 infection. (A and B) MHV68 latently infected A20-HE-RIT B cells, or parental A20 B cells, were treated with doxycycline and phorbol ester to induce lytic reactivation. Total RNA was isolated at the indicated time-points post-reactivation and subjected to primer extension for B2 SINEs or 7SK (as a loading control). (C) BMDMs or (D) NIH3T3 were either mock infected or infected with MHV68 for the indicated time periods, whereupon total RNA was isolated and subjected to primer extension as described above. The relative induction of B2 RNA was calculated by normalizing both to the 7SK loading control and to the levels of B2 RNA present in mock infected cells, which were set to 1.

We next examined the potential for B2 SINE induction upon MHV68 infection of primary bone marrow-derived macrophages (BMDMs). Unlike fibroblasts, which are highly susceptible to MHV68, the highest level of infection we achieved in WT BMDMs was ∼20%, which occurred with a multiplicity of infection (MOI) of 20, and did not increase upon addition of more virus (unpublished observation). Despite the lower infection efficiency, primer extension reactions demonstrated that in MHV68 infected primary BMDMs, B2 SINE induction began at 30 hpi and reached maximal levels by 40-48 hpi (Fig 1C). These induction kinetics were slower than what we observed in NIH 3T3 cells (Fig 1D), likely due to overall slower replication kinetics of MHV68 in the BMDMs. In summary, B2 SINE RNA induction occurs during lytic MHV68 infection of multiple primary and immortalized cell types.

### B2 SINE RNAs are not induced in uninfected cells by paracrine signaling

We were struck by the robust B2 upregulation in primary BMDMs, given that at most 20% of these cells were infected by MHV68. We therefore considered the possibility that infected cells produce paracrine signals that cause B2 upregulation in neighboring uninfected cells as well. We first tested this possibility using 3T3 cells, as their susceptibility to infection should yield a higher concentration of relevant paracrine signaling molecules. We performed a supernatant transfer assay, in which uninfected cells were incubated for 1 h or 24 h with cell supernatants from infected NIH 3T3 cells, either in crude form or after 0.1 μm filtration to remove viral particles. B2 SINE levels were then measured 24 h post transfer using primer extension. We observed no B2 SINE induction in cells incubated in filtered supernatants, suggesting that paracrine signals derived from infected 3T3 cells are not sufficient to stimulate B2 induction in uninfected cells. In contrast, there was robust B2 SINE induction in cells incubated with crude supernatants, as expected since these supernatants contain MHV68 virions to initiate a de novo infection (Fig 2A). This experiment was repeated in BMDMs, where filtered or crude supernatants were taken from infected 3T3 cells and incubated with plated BMDMs for 1 h or 24 h before removal. BMDMs were harvested 48 h after the beginning of incubation with 3T3 supernatants. These data were identical to that observed with 3T3s, in which paracrine signals contained within filtered supernatant were insufficient for B2 induction (Fig 2B).

**Figure 2:**
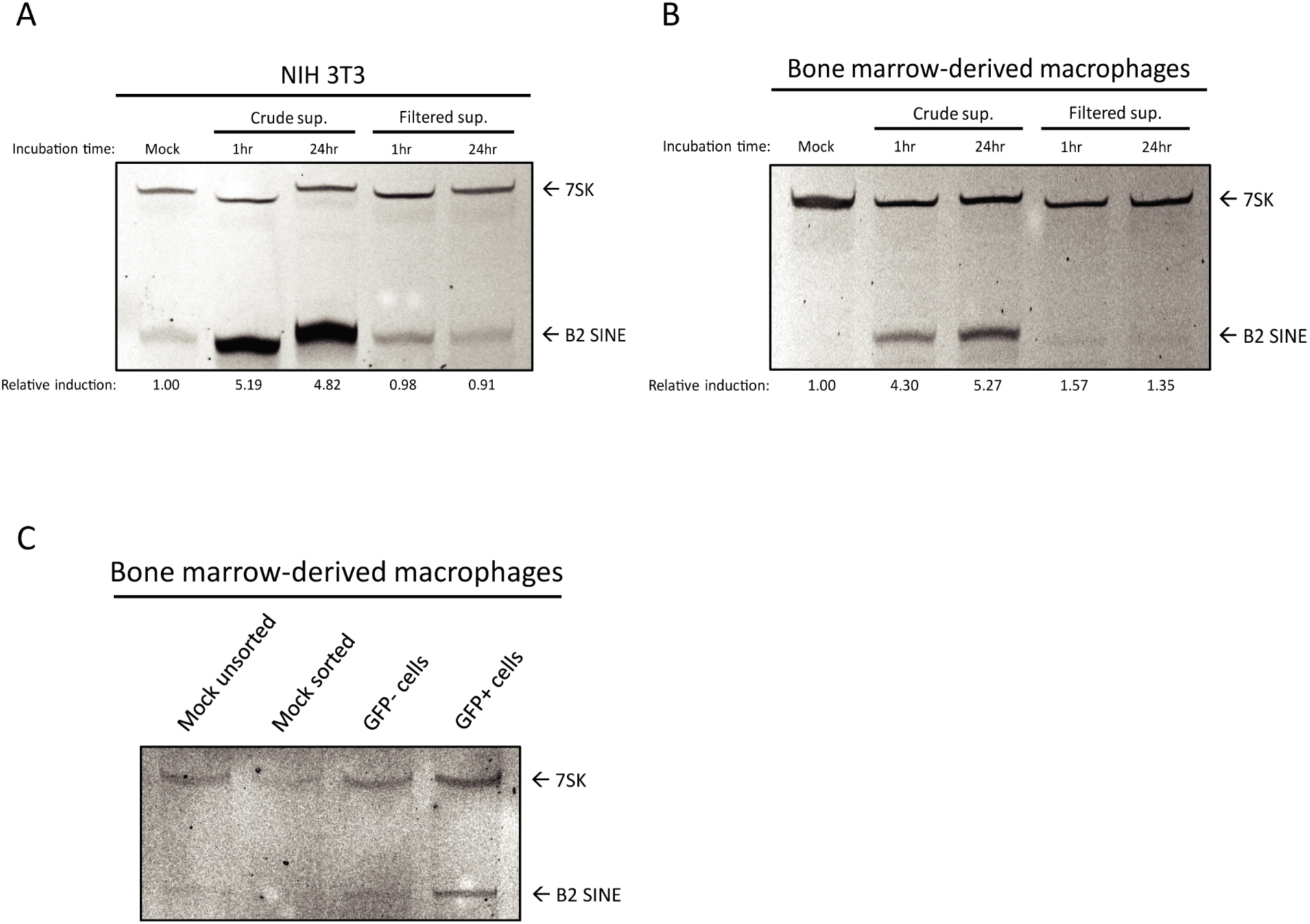
Paracrine signaling does not induce B2 SINE induction. (A) NIH3T3 cells or (B) primary BMDMs were incubated with supernatants harvested from 24 h infected NIH3T3 cells, either in crude form, or filtered to remove whole virus, for the indicated time period. Total RNA was isolated from cells at 24 h or 48 h post-incubation, respectively, and subjected to primer extension for B2 SINEs or 7SK. The relative induction of B2 RNA was calculated by normalizing both to the 7SK loading control and to the levels of B2 RNA present in mock infected cells, which were set to 1. (C) Infected BMDMs were sorted by flow cytometry to separate GFP+ (infected) from GFP-(uninfected) cell populations. Total RNA was isolated from each population and subjected to primer extension for B2 SINEs or 7SK.

We also looked for evidence of paracrine-based B2 induction in the primary BMDMs using a cell sorting strategy. Here, we made use of the fact that in a given infection assay, only 20% of the BMDMs will be infected with MHV68. Because we were using MHV68 containing a constitutively expressed GFP marker, we sorted GFP positive (infected) from GFP negative (uninfected) cells and performed B2 SINE primer extensions on each population (Fig 2C). As a control, we also sorted mock infected cells and confirmed that the stress of the sorting procedure did not activate B2 SINE transcription. We observed a greater B2 SINE signal in the GFP positive population, while the GFP negative population closely matched that of our negative uninfected control population. Together, these results suggest that induction of B2 SINE RNA occurs only in MHV68 infected cells and that paracrine or cell-to-cell signaling through the supernatant is not sufficient to induce this phenotype.

### B2 SINE induction is RNAPIII-dependent but does not involve the RNAPIII regulator Maf1

We previously showed that treatment of 3T3 cells with a RNAPIII inhibitor or B2-directed antisense oligonucleotides (ASOs) reduced the B2 RNA levels upon MHV68 infection, strongly suggesting that RNAPIII activity was required for their induction [21]. However, given that small molecule inhibitors can have off target effects and B2 ASOs will also target mRNAs containing embedded SINE elements, we sought to independently validate that the B2 SINE transcriptional induction is RNAPIII-dependent. We chose the strategy of depleting Brf1, a critical component of the TFIIIB transcription factor complex needed for RNAPIII transcription of type-II (e.g. SINE) promoters using siRNA-mediated knockdown [8]. Knockdown of Brf1 was robust through 48 h post-transfection (Fig 3A). In both BMDMs (Fig 3B) and 3T3 cells (Fig 3C), depletion of Brf1 completely abrogated B2 expression as measured by primer extension throughout the time course of infection. Notably, the levels of 7SK were not affected by Brf1 knockdown, as this RNAPIII transcript has a type III promoter that does not require Brf1 [40]. Thus, these results confirm that RNAPIII is required for MHV68-induced B2 SINE activation.

**Figure 3:**
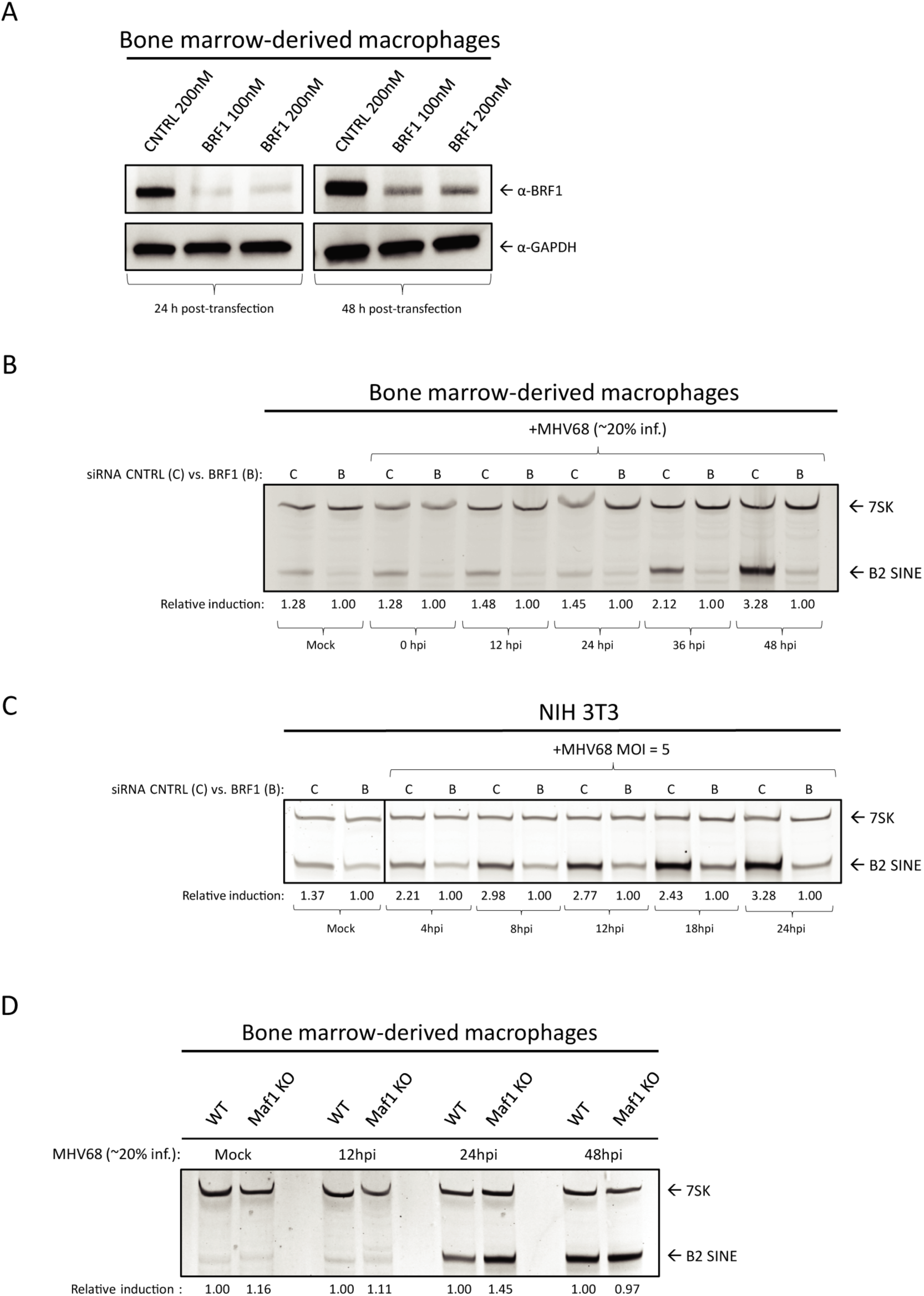
B2 SINE upregulation is dependent on RNAPIII, but independent of the RNAPIII master regulator Maf1. (A) BMDMs were transfected with the indicated concentrations of either control or Brf1 siRNA pools and harvested 24-48 h later. 30 μg of total protein lysates were resolved by SDS-PAGE and western blotted with antibodies against Brf1 or GAPDH (as a loading control). (B) Total RNA was harvested from mock or MHV68-infected BMDMs and NIH 3T3 fibroblasts following control or Brf1 siRNA treatment at the indicated time points. Total RNA was subjected to primer extension using primers for B2 SINEs or 7SK (as a control). (C) WT or (D) *Maf1-/-* BMDMs were mock infected or infected with MHV68 for the indicated times, whereupon total RNA was harvested and subjected to primer extension as described in (B). The relative induction of B2 RNA was calculated by normalizing both to the 7SK loading control and to the levels of B2 RNA present in Brf1 siRNA treated cells (B) or in WT BMDMs (C), which were set to 1.

We next considered the possibility that MHV68 infection alters the regulation of RNAPIII to increase its activity on B2 promoters. A master regulator of RNAPIII is Maf1, which acts by binding free RNAPIII at its clamp domain, thereby impairing RNAPIII binding to the TFIIIB-promoter complex and preventing RNAPIII transcription initiation [9, 41] To test the hypothesis that release of Maf1-mediated repression of RNAPIII transcription is responsible for B2 SINE induction, we derived primary BMDMs from *Maf1-/-* mice [42]. Surprisingly, we observed no increase in B2 SINE RNA in uninfected *Maf1-/-* BMDMs compared to WT BMDMs, suggesting that Maf1 is not required for the normal silencing of B2 loci (Fig 3D). We did observe somewhat more of an increase in B2 levels at 24 hpi with MHV68 in the *Maf1-/-* cells relative to WT cells, although this difference was not sustained at 48 hpi (Fig 3D). We therefore conclude that the primary mechanism of B2 induction by MHV68 is not through interference with the RNAPIII repressor Maf1.

### B2 SINE induction is independent of canonical innate immune signaling pathways

Due to their activation during herpesvirus infection and broadly acting signaling cascades, we considered that innate immune signaling may be involved upstream of B2 SINE induction. Pattern recognition receptors, namely the toll-like receptors 2, 3, 7, and 9, RIG-I-like receptors, and AIM2, can become activated during lytic herpesvirus infection [43–46]. To examine the possible upstream involvement of infection-induced innate immune signaling in the induction of B2 SINE transcription, we quantified B2 SINE levels in primary BMDMs derived from WT B6 mice versus mice lacking several canonical innate immune signaling pathways. These included mutants in toll-like receptor signaling (MyD88/*TRIF -/-*), cytoplasmic RNA recognition signaling (*MAVS -/-*), or type-I interferon (IFN) receptor mediated signaling through the type-I IFN receptor (*IFNAR -/-*) (Fig 4A), as well as cGAS/STING-mediated DNA sensing using the golden ticket (gt/gt) mutant [47], which contain a missense mutation in exon 6 of the mouse STING gene, rendering STING inactive (Fig 4B). In each case, primer extension experiments showed equivalent or greater B2 SINE RNA induction upon infection of the mutant BMDMs compared to the WT BMDMs. Thus, none of these innate immune components is individually required for SINE activation during MHV68 infection.

**Figure 4:**
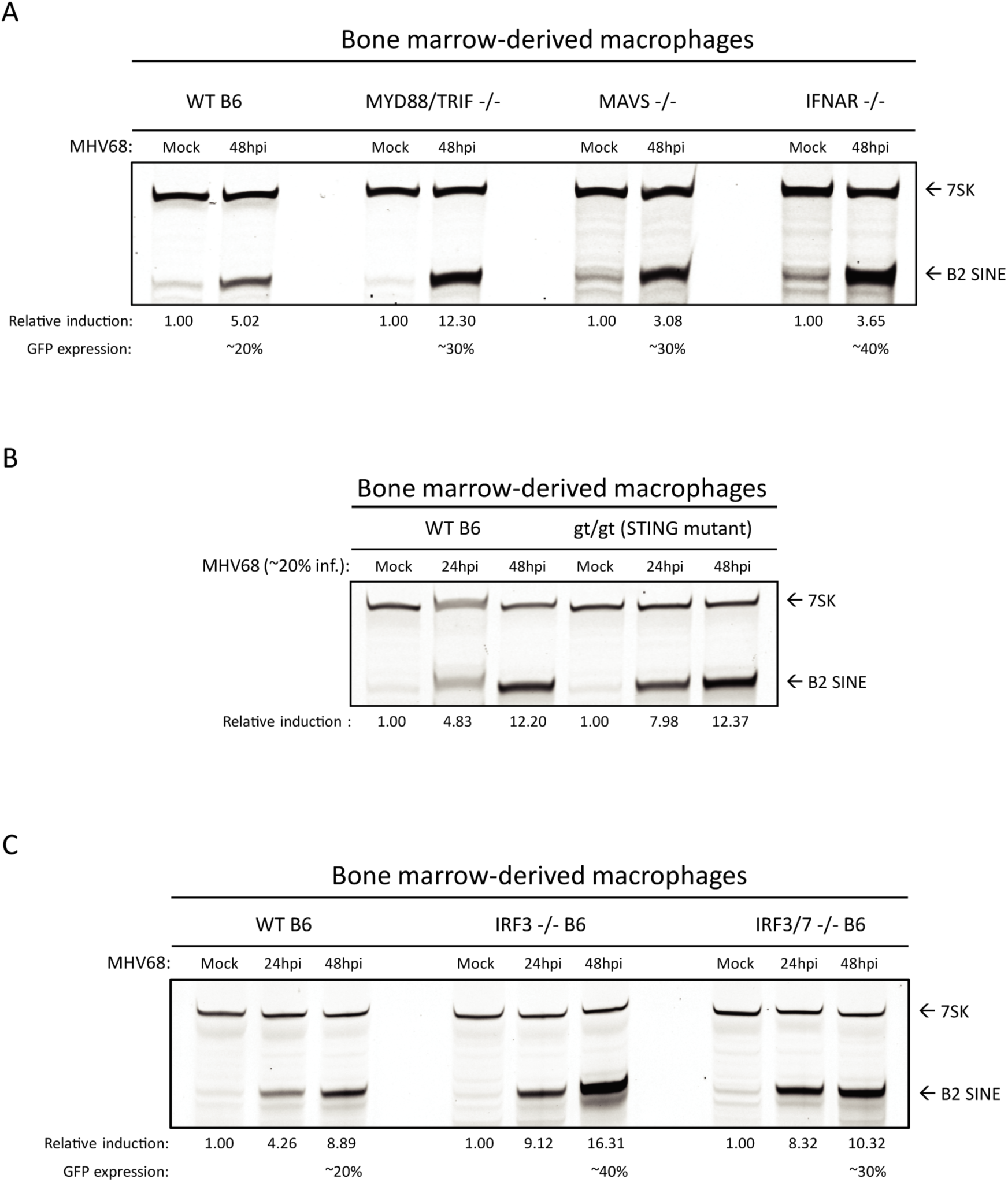
B2 SINE induction occurs independent of innate immune signaling. (A-C) WT or the indicated innate immune factor knockout BMDMs were mock or MHV68-infected for 24-48 h. Total RNA was then harvested and subjected to primer extension using primers for B2 SINEs or 7SK (as a control). The relative induction of B2 RNA was calculated by normalizing both to the 7SK loading control and to the levels of B2 RNA present in mock infected cells, which were set to 1.

To control for the possibility that multiple innate immune sensors could be activated in a redundant manner to induce B2 SINEs, we also tested primary BMDMs derived from mice lacking the downstream transcription factors interferon-regulatory factor 3 (IRF3) and interferon regulatory factor 7 (IRF7). All pattern recognition receptor signaling pathways converge on IRF3 and IRF7, which activate transcription of interferon stimulated genes (ISGs) and inflammatory cytokines [48]. In agreement with the data from BMDMs lacking the upstream innate immune sensors, MHV68 infection still caused robust B2 SINE induction in *IRF3 -/-* and *IRF3/7 -/-* BMDMs (Fig 4C). Thus, innate immune signaling does not activate B2 SINE transcription during MHV68 infection.

We noted that the infection-induced B2 levels were even more pronounced in each of the single and double knockout BMDMs than in WT cells (Figs. 4A-C). We hypothesize that this is a result of increased MHV68 infection under conditions of impaired immune restriction, as we noted that the knockout BMDMs routinely achieved higher MHV68 infection rates (as measured by GFP positivity) than WT BMDMs (unpublished observation).

### The conserved herpesvirus kinase ORF36 is sufficient to induce B2 SINE transcriptional upregulation

To search for viral factors involved in B2 SINE induction, we obtained and re-sequenced a partial MHV68 open reading frame (ORF) library previously generated by Dr. Ren Sun, which contained 47 full-length MHV68 ORF plasmids [49] (Table 1). The ORFs were first screened by co-transfection of 3T3 cells with 3-5 plasmids that were grouped based on similar temporal class and/or proposed or known function (Fig 5A, Table 1)[50, 51]. Only the group that contained ORFs 33, 35, and 36 showed B2 SINE induction above that of the control GFP expressing plasmid as measured by primer extension (Fig 5B). We then tested each of these ORFs individually for the ability to induce B2 SINEs, revealing that only MHV68 ORF36 expression was sufficient to upregulate B2 SINEs both as an untagged construct, as well as with an N-terminal FLAG-tag (Fig 5B).

**Figure 5:**
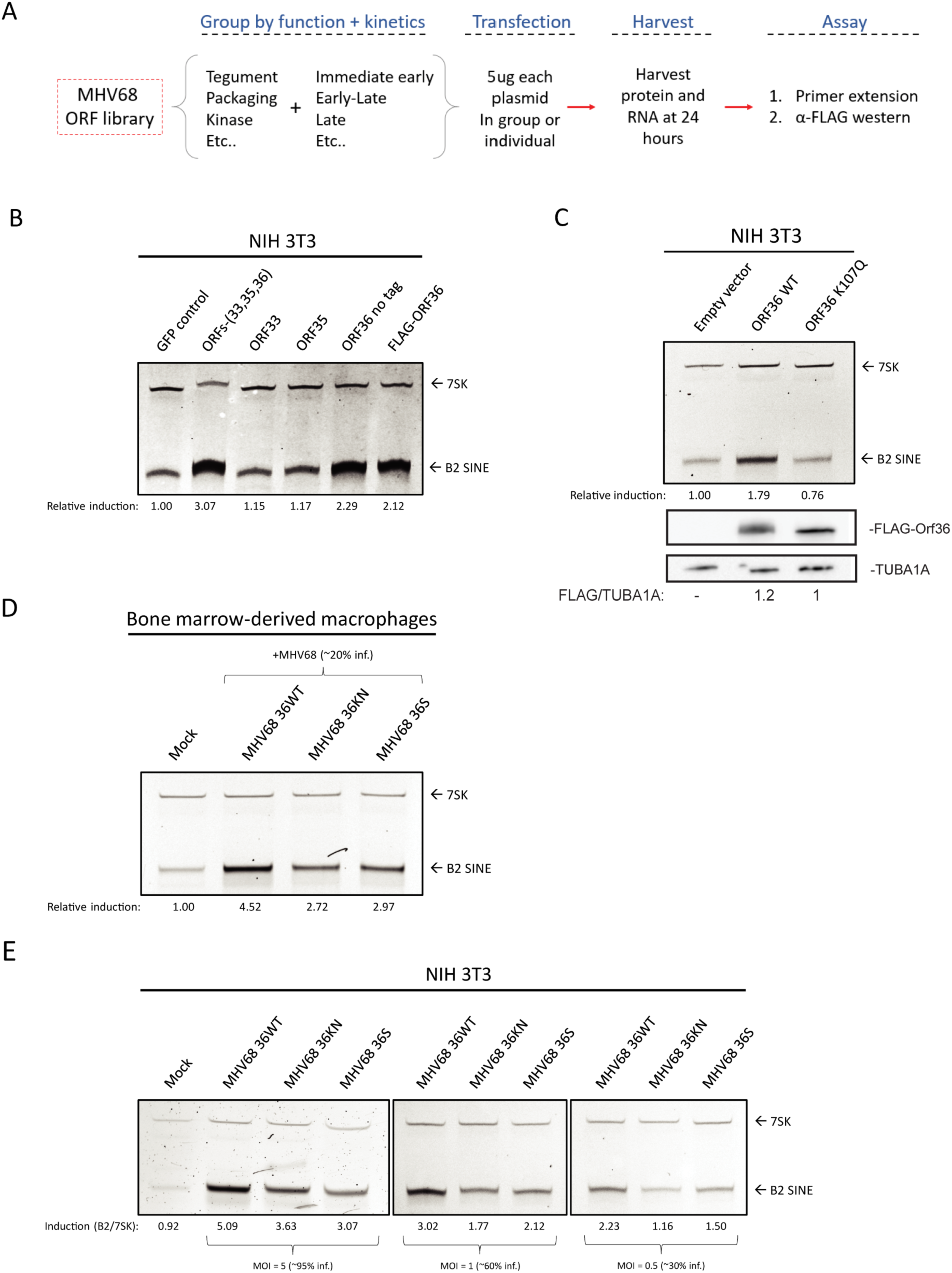
The MHV68 kinase ORF36 induces B2 SINE transcription. (A) Schematic representing the method for testing the MHV68 ORF library. (B) NIH3T3 cells were transfected with plasmid(s) containing the indicated ORF(s) or a GFP control for 24h, whereupon total RNA was extracted and subjected to primer extension using primers for B2 SINEs or 7SK (as a control). (C) NIH3T3 cells were transfected with plasmids expressing either wild-type (WT) ORF36 or a kinase null mutant (K107Q) for 24h then total RNA was isolated and subjected to primer extension as described above. (D) BMDMs were infected with WT MHV68, kinase null (KN), or ORF36 stop (S) virus at an MOI of 0.25. Total RNA was isolated at 48 hpi and subjected to primer extension as described in (B). (E) NIH 3T3 cells were infected with WT MHV68, KN, or S virus at an MOI of 5. At 24 hpi, total RNA was isolated and subjected to primer extension as described in (B). The relative induction of B2 RNA was calculated by normalizing both to the 7SK loading control (E) and to the levels of B2 RNA present in control transfected cells (B-C) or mock infected cells (D), which were set to 1.

**Table 1:** MHV68 ORFs tested in screen. ORFs were grouped (last column) based on similarities of kinetic class and function.

| ORF | Kinetic class | Proposed function | Group number |
| --- | --- | --- | --- |
| 49 | E-L | *unknown* | 1 |
| 50 | IE | Rta | 1 |
| 57 | IE | *unknown* | 1 |
| 73 | IE | LANA (transactivator) | 1 |
| 74 | E-L | GPCR | 1 |
| 40 | E-L | Helicase/primase | 2.1 |
| 60 | E-L | Ribonucleotide reductase | 2.1 |
| 61 | L | Ribonucleotide reductase | 2.1 |
| 56 | E-L | DNA repair helicase/primase | 2.2 |
| 59 | L | DNA repair/processivity factor | 2.2 |
| 7 | E-L | Tegument transport protein | 3 |
| 19 | L | Tegument (Thymidine kinase) | 3 |
| 32 | E-L | Tegument | 3 |
| 68 | E | Packaging | 3 |
| 11 | L | *unknown* | 4.1 |
| 20 | E-L | Fusion protein | 4.1 |
| 66 | L | Capsid | 4.1 |
| M1 | E-L | Unique secreted: interacts with ER | 4.2 |
| M3 | E-L | Unique secreted: chemokine binding | 4.2 |
| M11 | E-L | Bcl-2 homolog | 4.2 |
| 8 | L | gB | 5.1 |
| 22 | L | gH | 5.1 |
| 27 | L | Glycoprotein | 5.1 |
| 28 | L | Glycoprotein | 5.1 |
| 47 | E | gL | 5.2 |
| 53 | L | gM | 5.2 |
| M7 | L | gp150 | 5.2 |
| 10 | E-L | Inhibitor of mRNA transport | 6 |
| 37 | E-L | muSOX (alkaline exonuclease) | 6 |
| 33 | L | *unknown* | 7.1 |
| 35 | E-L | Tegument | 7.1 |
| 36 | IE | Tegument (Serine/Threonine Kinase | 7.1 |
| 42 | L | Tegument | 7.2 |
| 45 | L | *unknown* | 7.2 |
| 48 | L | *unknown* | 7.2 |
| 55 | E-L | *unknown* | 7.2 |
| 63 | E | Tegument | 7.2 |
| 23 | L | *unknown* | 7.3 |
| 38 | IE | Tegument | 7.3 |
| 58 | L | Membrane spanning protein | 7.3 |
| 24 | E | Late gene expression | 8 |
| 30 | E-L | Late gene expression | 8 |
| 31 | E-L | Late gene expression | 8 |
| 34 | E-L | Late gene expression | 8 |
| 72 | E-L | v-cyclin | 9 |
| M5 | E-L | *unknown* | 9 |
| M9 | E-L | *unknown* | 9 |

MHV68 ORF36 is a conserved herpesvirus serine/threonine kinase with a variety of reported kinase-dependent and -independent roles relating to the DNA damage response, inhibition of histone deacetylation, and inhibiting IRF3-driven ISG production [33, 34, 52–54]. To determine whether ORF36 kinase activity was required for B2 SINE upregulation, we compared the activity of WT ORF36 to an ORF36 kinase null mutant (K107Q) [54]. Primer extension of RNA from transfected 3T3 cells showed that only WT ORF36 but not K107Q induced B2 SINEs (Fig 5C). To determine the contribution of ORF36 towards B2 induction in the context of infection, we obtained versions of MHV68 either lacking ORF36 (ORF36 Stop (S)) or containing a kinase-null version of ORF36 (ORF36 KN) [53]. Notably, infection of primary BMDMs with these viruses revealed a reduction in MHV68-induced B2 SINE RNA upon loss or kinase inactivation of ORF36 compared to infection with the repaired WT virus (Fig 5D). We observed similar defects in B2 induction upon infection of 3T3 cells with ORF36 S and KN viruses compared to WT, across a range of MOI (Fig 5E). The fact that some residual B2 induction remained in BMDM and 3T3 cells infected with the ORF36 mutant viruses indicates that other viral factors also contribute to SINE induction. However, ORF36 expression is sufficient to activate B2 SINEs when expressed alone, and is required for WT levels of B2 SINE induction in the context of MHV68 infection.

### Induction of B2 SINE transcription is conserved amongst ORF36 CHPK homologs

ORF36 homologs are found in all subfamilies of herpesviruses, where they are collectively referred to as the Conserved Herpesvirus Protein Kinases (CHPKs). Several examples exist of shared CHPK functions and shared substrate specificity [55, 53, 56]. We therefore examined whether other CHPKs were able to induce B2 SINE RNA. We transfected NIH 3T3 cells with plasmids expressing HA- or FLAG-tagged CHPKs from KSHV (ORF36), varicella zoster virus (VZV) (ORF47), human cytomegalovirus (HCMV)(UL97), EBV (BGLF4), and MHV68 (ORF36) and measured B2 SINE RNA using primer extension (Fig 6A). MHV68 ORF36 produced the most robust induction, followed by the other gammaherpesvirus CHPKs, KSHV ORF36 and EBV BGLF4. The alpha- and betaherpesvirus protein kinases, VZV ORF47 and HCMV UL97, induced B2 SINEs to a minimal degree, although they were expressed to similar (albeit low) levels as MHV68 ORF36 (Fig 6B). Thus, while the ability to induce B2 SINE RNA appears to be conserved amongst the CHPKs, this function is most prominent among the gammaherpesvirus homologs.

**Figure 6:**
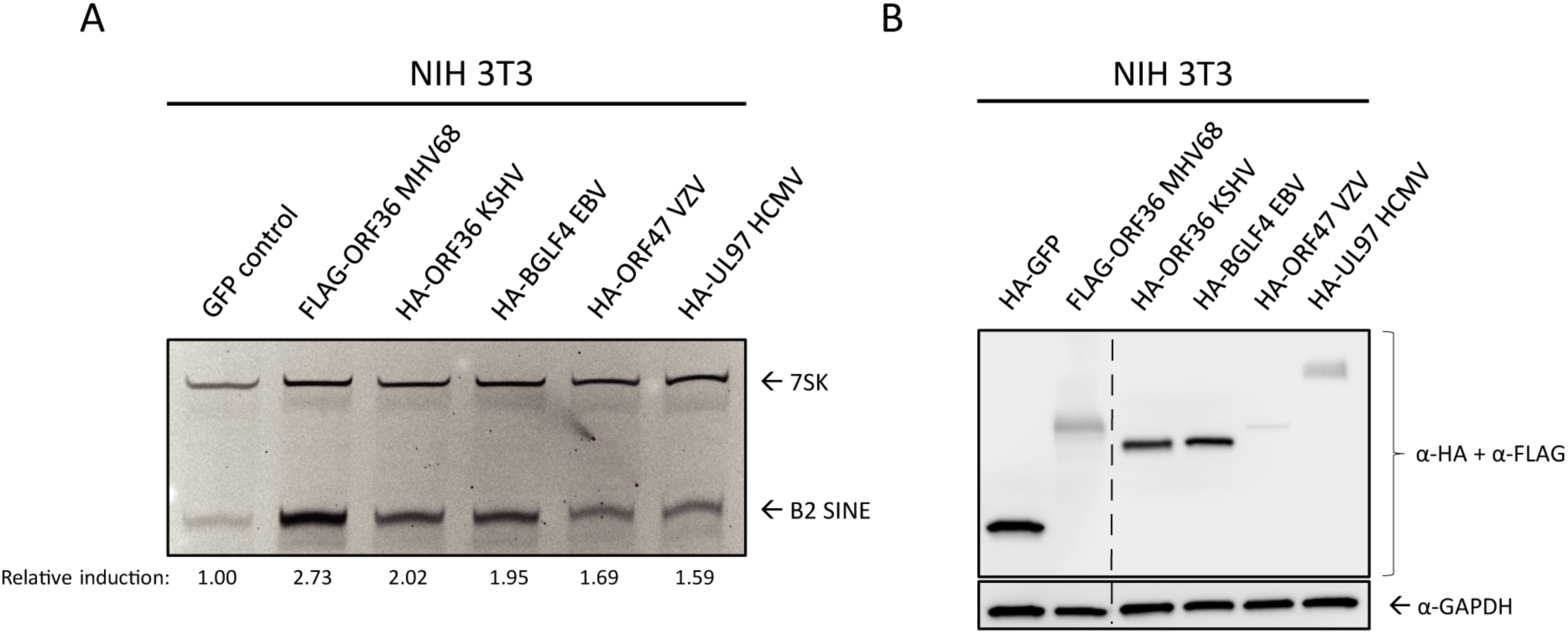
Functional conservation of B2 SINE upregulation by several MHV68 ORF36 homologs. NIH3T3 cells were transfected with plasmids containing FLAG-tagged MHV68 ORF36 or the indicated HA-tagged ORF36 homolog from Kaposi’s sarcoma-associated herpesvirus (KSHV ORF36), Epstein-Barr virus (EBV BGLF4), varicella zoster virus (VZV ORF47), or human cytomegalovirus (HCMV UL97). These cells were then harvested for total RNA for B2 and 7SK primer extension (A), or protein lysates, which western blotted with antibodies against HA and FLAG, or GAPDH as a loading control (B). The relative induction of B2 RNA was calculated by normalizing both to the 7SK loading control and to the levels of B2 RNA present in GFP control transfected cells, which were set to 1.

### De-repression of the chromatin landscape allows for B2 SINE induction

Previous studies of features linked to SINE repression in uninfected cells indicated the importance of the repressive histone H3 lysine 9 tri-methylation (H3K9me3) mark and, to a lesser degree, DNA methylation at CpG sites [11, 12, 57, 58]. These marks are deposited and maintained by the histone methyltransferases SU(VAR)3-9 and the DNMT family of DNA methyltransferases, respectively. Furthermore, ORF36 has been shown to inhibit histone deacetylases 1 and 2 (HDACs 1/2) [33], although whether HDACs are involved in repression of SINE loci is unknown.

To test the role of each of these factors in B2 induction, we treated NIH 3T3 cells with inhibitors of HDACs 1/2 (ACY-957), DNMTs (5-azacytidine), and SU(VAR)3-9 (chaetocin), or a cocktail composed of ACY-957 and chaetocin together (Fig 7A). We observed induction of B2 SINEs following treatment with ACY-957 and chaetocin, and an additive effect when using both inhibitors together (Fig 7A, lane 5). Treatment of cells with 5-azacytidine yielded no increase in levels of B2 RNA, in agreement with previous work [11].

**Figure 7:**
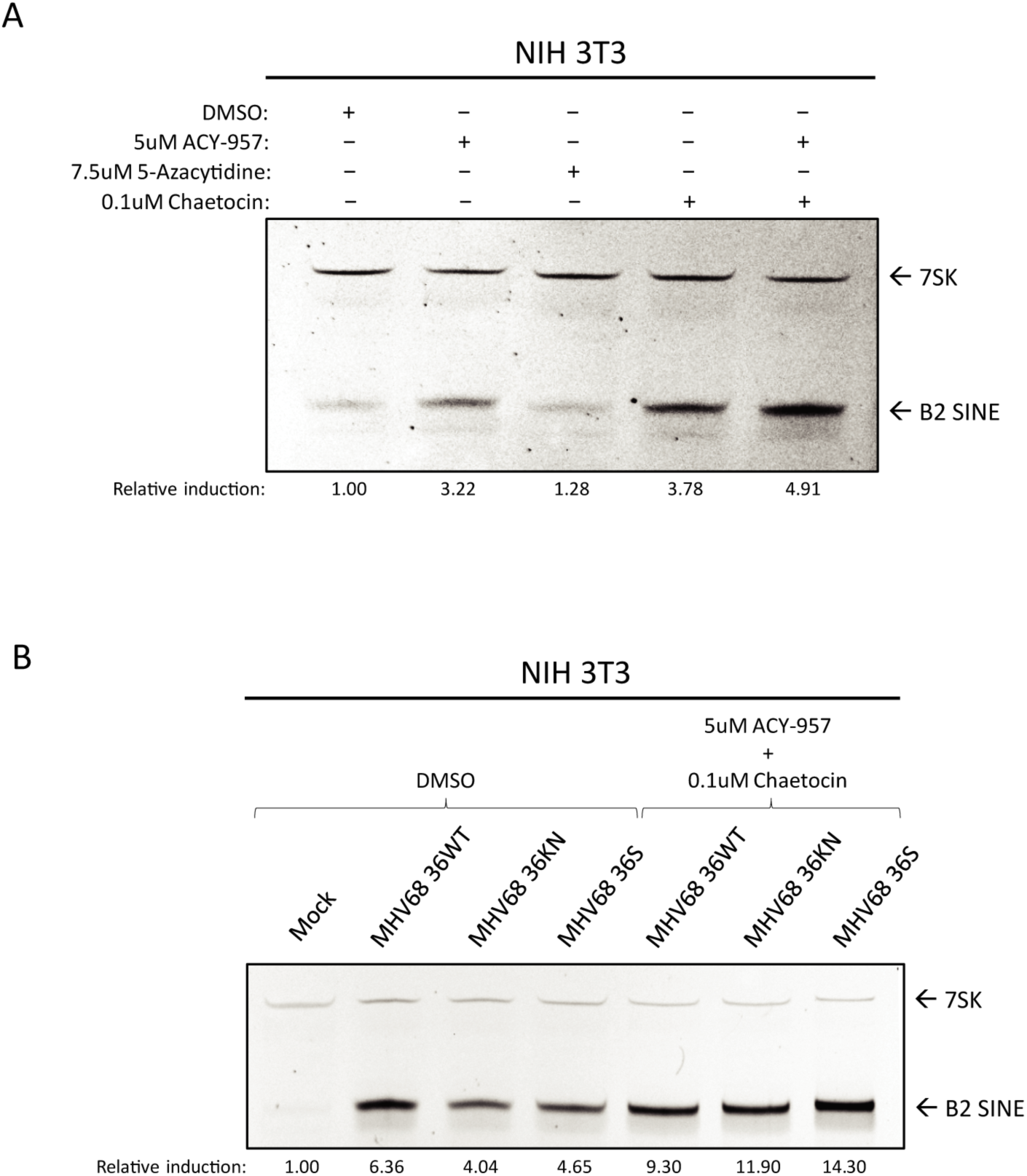
Inhibitors of chromatin repression cause B2 SINE upregulation. (A) NIH3T3 cells were treated with the indicated inhibitor(s) for 24 h, whereupon total RNA was isolated and subjected to primer extension for B2 SINEs or 7SK. (B) NIH3T3 cells were subjected to pre-treatment with DMSO or the indicated inhibitors for 1 h prior to infection with MHV68 WT, KN, or S virus for 24 h, whereupon total RNA was isolated and subjected to primer extension as described in (A). The relative induction of B2 RNA was calculated by normalizing both to the 7SK loading control and to the levels of B2 RNA present in DMSO treated (A) or mock infected cells (B), which were set to 1.

Given that the strongest effects on B2 induction were observed upon inhibition of histone methyltransferases combined with HDAC inhibition, we next tested whether treatment with these inhibitors during infection was sufficient to rescue B2 levels in ORF36 KN and S infection to ORF36 WT infection levels. We observed that, in the context of infection, treatment with ACY-957 and chaetocin restored the levels of B2 ncRNA in the ORF36 S and KN infected cells to those observed during WT MHV68 infection (Fig 7B), showing that chromatin de-repression induced B2 ncRNA accumulation in an additive manner. Taken together, these data show that keeping an actively repressed chromatin state, primarily through maintenance of H3K9me3, is important for preventing constitutive B2 SINE induction.

## DISCUSSION

A growing body of literature indicates that RNAPIII transcripts are upregulated in herpesvirus-infected cells and can serve as substrates for innate immune recognition, although mechanisms underlying their induction remain largely unknown [24, 59–61]. The most robustly induced class of such transcripts in MHV68 infected fibroblasts are the B2 SINE ncRNAs, whose transcription becomes activated across tens of thousands of loci [22]. Here, we show that B2 SINEs are also strongly induced in an RNAPIII-dependent manner in reactivated B cells and primary bone marrow derived macrophages, confirming that B2 activation is a prominent feature of MHV68 infection in physiologically relevant cell types. Induction of B2 SINEs occurs in a cell autonomous manner and they are not activated in uninfected cells via paracrine signaling. Furthermore, our data suggest that B2 induction is not a downstream product of antiviral signaling upon MHV68 infection, nor does it involve Maf1, a key negative regulator of RNAPIII activity. Instead, we link B2 activation to the conserved herpesviral serine/threonine protein kinase ORF36, which is sufficient to activate B2 RNA on its own and contributes to robust B2 accumulation during MHV68 infection. We hypothesize that changes in chromatin modification contribute to ORF36-mediated B2 activation, and that this activity is at least partially conserved in other herpesviral protein kinase homologs.

Several immune sensing pathways can become activated during lytic herpesvirus infection, and B2 induction in uninfected cells has been linked to various types of cell stress. TLRs 2, 3, and 9, as well as the DNA sensing AIM-2 like receptor family, the MAVS-dependent RNA-recognition receptors Mda5 and RIG-I, and the type-I interferon signaling pathway have all been implicated in the sensing of herpesviral infection [44, 46, 61–63]. However, our data from a variety of pattern recognition receptor and pathway knockout BMDMs indicate that engagement of these innate immune signaling components is not the mechanism by which MHV68 infection activates B2 SINEs. Indeed, B2 induction is even more robust in these infected knockout cells compared to WT BMDMs, likely reflecting enhanced replication of the virus in the absence of intact antiviral signaling. The innate immune-independence of B2 activation is in agreement with the timing of B2 induction, which initiates with delayed early kinetics and continually increases late in infection.

RNAPIII transcription is broadly impacted by Maf1, which binds and negatively regulates polymerase activity [9]. Thus, if B2 SINE induction were due to inactivation of Maf1, then we anticipated that *Maf1-/-* cells would have high baseline levels of B2 SINE RNA that would not further increase upon MHV68 infection. However, we did not observe any increase in B2 SINE levels in mock-infected *Maf1-/-* cells and MHV68 infection of these cells resulted in B2 SINE activation that was comparable to WT cells. These findings indicate that regulation of Maf1 does not influence MVH68-mediated B2 SINE activation. Consistent with this, a recent chromatin immunoprecipitation-sequencing study of RNAPIII occupancy in wild-type mouse liver found relatively few B2 SINEs and identified only ∼30 of these elements with increased RNAPIII occupancy in *Maf1-/-* mouse liver [64]. We did observe a slight increase in B2 SINE levels at 24 hpi in *Maf1-/-* compared to WT cells, suggesting quicker RNAPIII transcription kinetics due to broad loss of Maf1-mediated repression (Fig 3D).

A partial MHV68 ORF library screen revealed ORF36 to be a robust inducer of B2 SINE transcription. ORF36 is an early transcript [50, 51], which is consistent with the kinetics of B2 induction and with our current and prior observations that inhibition of viral DNA replication and late gene expression does not block B2 activation [21]. Like other CHPKs, ORF36 displays homology to the host-encoded cyclin-dependent kinases but is thought to have broader substrate specificity [56]. Indeed, it has been reported to phosphorylate many targets, including the retinoblastoma protein, H2AX, and lamin A/C [55, 65]. Additionally, ORF36 has kinase-independent functions such as inhibition of HDACs 1/2 [34] and IRF-3 [54], both of which are beneficial for productive infection. Given our results showing that pharmacological inhibition of HDACs 1/2 and SU(VAR)3-9 stimulated B2 induction, we favor the hypothesis that ORF36 activities related to chromatin remodeling underlie its B2 induction phenotype. This would be in line with previous work in uninfected cells demonstrating that DNA CpG methylation and histone H3 trimethylation (H3K9me3) contribute to transcriptional repression of SINE loci [11, 12, 57, 58]. The observation that the ORF36 kinase null viral mutant was as defective as the ORF36 stop mutant for B2 induction indicates that while ORF36 modulation of HDACs 1/2 may contribute to such chromatin remodeling, this kinase-independent function of ORF36 is not the primary driver of B2 induction during infection. Instead, it may facilitate sustained B2 activation following a kinase-dependent initial activation event.

Whether ORF36 impacts SU(VAR)3-9 methyltransferases is unknown, although phospho-proteomics analysis of the EBV CHPK, BGLF4, suggests that SU(VAR)3-9h2 is phosphorylated in a BGLF4-dependent manner [66]. An intriguing possibility is that ORF36 inhibits SU(VAR)3-9 function, either through direct phosphorylation of SU(VAR)3-9 or manipulation of an upstream regulator such as its repressor DBC1 [67]. Additionally, recruitment of heterochromatin protein 1 (HP1) to H3K9me3 marks is dependent on HDAC activity [68], providing another link between these chromatin regulatory factors. Future experiments will be geared towards exploring epigenetic alterations to the host genome during MHV68 infection that could influence RNAPIII transcription.

The viral protein kinases are emerging as important players in gammaherpesvirus-associated lymphomagenesis, and an intriguing possibility is that its activation of Pol III retrotransposons—which are known to cause insertional mutagenesis [69–71] — may contribute to this phenotype. Indeed, prolonged expression of the ORF36 homolog in EBV (BGLF4) can contribute to genome instability leading to tumor formation, which has been linked to its phosphorylation of lamin A/C and topoisomerase-II [72, 73]. KSHV ORF36 also displays functions associated with oncogenesis, including functional mimicry of the cellular ribosomal protein S6 kinase β-1 (S6KB1), which leads to enhanced protein synthesis, endothelial capillary tubule formation and anchorage-independent growth [74]. Notably, a recent study from the Damania lab showed that transgenic mice expressing KSHV ORF36 display increased B cell activation and develop high-grade B cell lymphomas that share many features of primary effusion lymphoma [75]. In this regard, it is notable that among the vPK homologs, EBV BGLF4 and KSHV ORF36 showed the highest degree of B2 activation. The extent to which Pol III activation contributes to these oncogenic phenotypes, as well as whether MHV68 ORF36 also contributes to lymphomagenesis are important questions for the future.

MHV68 viral mutants lacking ORF36 or expressing a kinase null version of the protein displayed a partial reduction in B2 RNA accumulation relative to WT virus. These results suggest that while ORF36 contributes to B2 induction during infection, one or more other viral activities may be involved. Our ORF screen encompassed a significant percentage of the annotated MHV68 genome [76], however it should be noted that recent work from O’Grady et al. [77] shows pervasive alternate isoform usage overlapping ORF isoforms, suggesting that MHV68 encodes a more diverse proteome than previously anticipated. One or more of these untested proteins may also contribute to B2 induction, either via independent mechanisms or in cooperation with ORF36. Investigations of other MHV68-encoded ORFs involved in B2 SINE transcription and stabilization remains an open area of investigation.

In summary, our results provide the first insights into how gammaherpesvirus infection induces SINE retrotransposons, and identify a novel activity of the ORF36 protein kinase. Our work supports a model in which ORF36 kinase-dependent and –independent functions inhibit proteins involved in the maintenance of a repressive chromatin landscape, thereby contributing to de-repression of B2 SINEs. How these activities selectively impact certain RNAPIII loci remains a key open question. Indeed, ongoing work to define how SINEs and other RNAPIII transcripts are activated during infection, as well as noncanonical functions of these ncRNAs, should provide insight into the emerging field of retrotransposon-linked cell signaling. Given the breadth of DNA viruses that activate these hyper-abundant loci, viruses will continue to serve as unique tools to dissect the regulation of ncRNAs, as well as the mechanisms by which they influence the outcome of infection.

## MATERIAL and METHODS

### Cells

NIH 3T3 mouse fibroblasts were obtained from the UC Berkeley Cell Culture Facility and maintained in Dulbecco’s modified Eagle’s Medium (DMEM; Invitrogen) with 10% fetal calf serum (FBS; Seradigm). A20 B cells (kindly provided by Laurie Krug, [39]) were maintained in RPMI (Gibco), 10% fetal bovine serum (FBS; VWR), 2 mM L-glutamin, 100 U/ml penicillin, 100 mg/ml streptomycin, and 50 mM BME. A20-HE-RIT cell lines (kindly provided by Laurie Krug, [39]) were maintained under the same conditions as A20 cells, with the addition of 300 µg/ml hygromycin B, 300 µg/ml G418 and 2 µg/ml puromycin. To reactivate A20-HE-RIT, cells were cultured in media without antibiotic selection for 24 h, then seeded at a cell density of 1.0X10^6^ cells/ml in the presence of 5 µg/ml doxycycline and 20 ng/µl PMA for the indicated time. To block viral DNA replication, PAA was used at a concentration of 200 μg/ml and was added at the start of reactivation. Bone marrow-derived macrophages (BMDMs) containing knockouts for innate immune pathway components [47, 78–80] were kindly provided by the lab of Dr. Gregory Barton (UC Berkeley, Department of Immunology). Wild-type and *Maf1* knockout BMDMs were differentiated as follows: Femurs and tibias from C57BL/6J (B6) mice [42] aged 3-6 months were flushed with bone marrow media + antibiotics (BMM+A; High glucose DMEM + 10%FBS, + 10% MCSF + 1%PenStrep) using a 3 mL syringe with attached 23-gauge needle. Cell-containing media was filtered through a 70 μM filter to remove debris. Cells were pelleted at 280 x *g* in an Allegra X-15R Beckman Coulter centrifuge for 5 minutes. Supernatant was removed by aspiration and cells resuspended in BMM+A. Cells were counted using a hemocytometer and plated in non-TC treated 15CM petri dishes (Falcon, Ref #351058) at a concentration of 10e6 cells/25mL BMM+A/plate. On day 3 of differentiation, 5 mL BMM+A was added to each plate to feed cells. On day 7 of differentiation, BMM+A was aspirated and replaced with 10 mL cold Dulbecco’s Phosphate-Buffered Saline (DPBS; Invitrogen) per plate and placed at 4°C for 10 min. Cells were then lightly scraped from each plate and collected, pelleted as previously mentioned, and resuspended in bone marrow media without antibiotics (BMM) containing 10% DMSO at a concentration of 10e6/mL. 1.5 mL CryoTube^™^ Vials containing 1 mL/10e6 BMDMs were frozen at −80°C for 24 h before being stored in liquid nitrogen for duration. Subsequently, thawed vials of BMDMs were maintained in BMM except during infections.

### Plasmids and cloning

MHV68 ORF library plasmids were generously provided by the lab of Ren Sun (University of California Los Angeles) and their construction is previously described [49]. For generation of the ORF36 kinase-null mutant, the K107Q mutation was introduced by QuickChange PCR with the following primers: 5’-GTGCTGTCAATTTTGGGATATACTGTATGCAGAGCGTGTCATCTGAT-3’ and 5’-ATCAGATGACACGCTCTGCATACAGTATATCCCAAAATTGACAGCAC-3’. Plasmids for conserved herpesvirus protein kinase homologs of ORF36 were purchased through Addgene from the laboratory of Dr. Robert Kalejta (https://www.addgene.org/Robert_Kalejta/) [55].

### Virus preparation and infections

MHV68 containing a stop mutation or kinase null mutation in ORF36, as well as the corresponding mutant rescue virus, were generously provided by Vera Tarakanova (Medical College of Wisconsin) [53]. MHV68 was amplified in NIH 3T12 fibroblast cells, and the viral TCID50 was measured on NIH 3T3 fibroblasts by limiting dilution. NIH 3T3 fibroblasts were infected at the indicated multiplicity of infection (MOI) by adding the required volume of virus to cells in 1 mL total volume (for each well of a 6-well plate) 2 mL total volume (for 6cm plates) or 5 mL (for 10cm plates). Infection was allowed to proceed for 45 min prior to removal of virus media and replacement with DMEM + 10% FBS. BMDMs were infected with the minimal volume of MHV68 required to achieve maximum infection (20-30%), as determined by titration experiments with GFP-marked MHV68 followed by flow cytometry for GFP. For infection of BMDMs, virus was added to cells in serum-free DMEM for 4 h in non-TC treated plates. Virus containing media was then aspirated and replaced with macrophage media without antibiotics.

### Primer extension

Total RNA was extracted from cells using TRIzol reagent (Invitrogen). Primer extension was performed on 10-15 μg of total RNA using a 5’ fluorescein labeled oligo specific for B2 SINEs or 7SK. RNA was ethanol precipitated in 1 mL 100% EtOH, washed in 70% EtOH and pelleted at 21,130 x *g* and 4 °C for 10 min. Pellets were re-suspended in 9 μL annealing buffer (10 mM Tris-HCl, pH7.5, 0.3 M KCl, 1 mM EDTA) containing 1 μL of (10pmol/uL) 5’-fluorescein labeled primer (B2 SINE: TACACTGTAGCTGTCTTCAGACA; 7SK: GAGCTTGTTTGGAGGTTCT; Integrated DNA Technologies). Samples were heated briefly to 95 °C for 2 min, followed by annealing for 1 h at 55 °C. 40 μL of extension buffer (10 mM Tris-HCl, pH 8.8, 5 mM MgCl2, 5 mM DTT, 1 mM dNTP) and 1 μl of AMV reverse transcriptase (Promega) was then added and extension was carried out for 1 h at 42 °C. Samples were EtOH precipitated, then pellets were briefly air dried and resuspended in 20 μL 1X RNA loading dye (47.5% formamide, 0.01% SDS, 0.01% bromophenol blue, 0.005% xylene cyanol and 0.5 mM EDTA.). 10 μL of each sample was run on an 8% UREA-PAGE gel for 1 h at 250V. Gels were imaged on a Biorad Chemidoc with Fluoroscein imaging capability. Band intensity was quantified in ImageJ and normalized as described in the figure legends.

### Cell Sorting

For GFP expression of fixed cells: Cells were treated with 100 μL of trypsin for several min in well before being neutralized with 100 μL cold DPBS and transferred to a 96-well V-bottom plate in 200 μL total. They were then centrifuged at (475 x *g*) for 1 min. Media was removed and replaced in each well with 200 μL cold DPBS before being spun down again to wash. This was repeated twice. Cells in each well were then resuspended in 200 μL of 10% formaldehyde in DPBS to fix cells for 10 min at 4°C. The plate was then spun down and washed twice as described above. Cells were then resuspended in a final volume of 200 μL DPBS for cell sorting with a BD Accuri™ C6 Flow Cytometer.

For sorting of un-fixed GFP expressing cells, plates containing MHV68 infected BMDMs were washed twice with cold DPBS and cells were gently scraped from plates. Cells were centrifuged for 5 minutes at 475 x *g* to pellet, and then resuspended in warm BMDM media at a concentration of 5e6 cells/mL. Cell-containing media was passed through a 70μm filter into a 15mL conical. GFP+ and GFP-cells were sorted directly into TRIzol reagent using an Aria Fusion cell sorter.

### Protein extraction and analysis

Cells were washed with cold DPBS once before being lysed with RIPA lysis buffer (50 mM Tris HCl, 150 mM NaCl, 1.0% (v/v) NP-40, 0.5% (w/v) sodium deoxycholate, 1.0 mM EDTA, and 0.1% (w/v) SDS). Cell lysates were vortexed briefly, rotated at 4°C for 1 h, and then spun at 18,000 x *g* in a table-top centrifuge at 4°C for 12 min to remove debris.

For western blot analyses, 30 μg of whole cell lysate was resolved with 4-15% Mini-PROTEAN TGX gels (Bio-Rad). Transfers to PVDF membranes were done with the Trans-Blot Turbo Transfer system (Bio-Rad). Blots were incubated in 5% milk/TBS+0.1% Tween-20 (TBST) to block, followed incubation with primary antibodies against FLAG (Sigma F1804, 1:1000), Brf1 (Bethyl a301-228a, 1:1000), HA (Sigma H9658, 1:1000), TUBA1A (abcam ab729, 1:1000), or GAPDH (Abcam ab8245, 1:1000) in 5% milk/TBST. Washes were carried out in in TBST. Blots were then incubated with HRP-conjugated secondary antibodies (Southern Biotechnology, 1:5000). Washed blots were incubated with Clarity Western ECL Substrate (Rio-Rad) for 5 min and visualized with a Bio-Rad ChemiDoc.

### Inhibitor treatment

Cells were plated 12 h before inhibitor treatment to achieve 70% confluency at the time of treatment. ACY-957 (MedChemExpress HY-104008), 5-azacytidine (Sigma A2385), and chaetocin (Cayman Chemicals 13156), were re-suspended with DMSO prior to treatment. Inhibitors were diluted to working concentrations in warmed DMEM + 10% FBS before addition to cells. Pre-treatment of cells with inhibitor containing media preceded infection with MHV68 by 1 h. Upon removal of virus containing media, inhibitor containing media was replaced onto cells.

### Ethics Statement

All experiments involving mice were performed in accordance with the National Institute of Health’s Office of Laboratory Animal Welfare using protocols (20160305 and 20160311) approved by the Institutional Animal Care and Use Committee (IACUC) of the Albert Einstein College of Medicine, which fully accredited by the Association for the Assessment and Accreditation of Laboratory Animal Care (AAALAC).

## ACKNOWLEDGEMENTS

We wish to thank Ren Sun and Ting-Ting Wu (University of California Los Angeles) for providing their MHV68 ORF library, Vera Tarakanova (Medical College of Wisconsin) for providing the ORF36.stop and kinase null MHV68 and for helpful comments on the manuscript, Laurie Krug (Stony Brook University) for providing the A20-HE-RIT and A20 cells, and to the labs of Greg Barton and Russell Vance (University of California Berkeley) for sharing immune factor knockout macrophages. This work was funded by National Institutes of Health grants AI147183 and CA136367 to BG and GM120358 to IW. AS was funded by a graduate research fellowship from the National Science Foundation (DGE 1752814). BG is an investigator of the Howard Hughes Medical Institute.

